# RIG-I-induced innate antiviral immunity protects mice from lethal SARS-CoV-2 infection

**DOI:** 10.1101/2021.08.06.455405

**Authors:** Samira Marx, Beate M. Kümmerer, Christian Grützner, Hiroki Kato, Martin Schlee, Eva Bartok, Marcel Renn, Gunther Hartmann

**Affiliations:** Institute of Clinical Chemistry and Clinical Pharmacology, University Hospital Bonn, 53127 Bonn, Germany; Institute of Virology, University Hospital Bonn, 53127 Bonn, Germany; Institute of Cardiovascular Immunology, University Hospital Bonn, 53127 Bonn, Germany; Unit of Experimental Immunology, Department of Biomedical Sciences, Institute of Tropical Medicine, 2000 Antwerp, Belgium; Mildred Scheel School of Oncology, University Hospital Bonn, 53127 Bonn, Germany

## Abstract

The SARS-CoV-2 pandemic has underscored the need for rapidly employable prophylactic and antiviral treatments against emerging viruses. Nucleic acid agonists of the innate immune system can be administered to activate an effective antiviral program for prophylaxis in exposed populations, a measure of particular relevance for SARS-CoV-2 infection due to its efficient evasion of the host antiviral response. In this study, we utilized the K18-hACE2 mouse model of COVID-19 to examine whether prophylactic activation of the antiviral receptor RIG-I protects mice from SARS-CoV-2 infection. Systemic treatment of mice with a specific RIG-I ligand one to seven days prior to infection with a lethal dose of SARS-CoV-2 improved their survival of by up to 50 %. Improved survival was associated with lower viral load in oropharyngeal swabs and in the lungs and brain of RIG-I-treated mice. Moreover, despite antiviral protection, the surviving mice that were treated with RIG-I ligand developed adaptive SARS-CoV-2-specific immunity. These results reveal that prophylactic RIG-I activation by synthetic RNA oligonucleotides is a promising strategy to convey short-term, unspecific antiviral protection against SARS-CoV-2 infection and may be a suitable broad-spectrum approach to constraining the spread of newly emerging viruses until virus-specific therapies and vaccines become available.

## Introduction

The severe acute respiratory syndrome coronavirus 2 (SARS-CoV-2) pandemic has called to attention the vital importance of rapid, effective strategies for limiting the spread of emerging viruses. SARS-CoV-2 is the etiological agent of Coronavirus Disease 2019 (COVID-19)^1,2^, which infects the upper and lower airways of patients but also can cause neurological symptoms, in particular anosmia^3,4^. The clinical course of COVID-19 is extremely variable between individuals, from mild symptoms to severe interstitial pneumonia requiring mechanical ventilation. Since its initial outbreak in 2019 in Wuhan, SARS-CoV-2 virus infection has resulted in over 198 million confirmed COVID-19 cases and over 4.2 million deaths (https://COVID19.who.int/, Status: 2 August 2021). Moreover, we are only beginning to understand the extent of its socioeconomic repercussions, including the impact of chronic disease^5^, loss of primary caregivers ^6^, unemployment and school closures. While tremendous efforts are being made to control the virus, the development of vaccines has progressed more rapidly than the development of direct antiviral treatments^7^. Repurposed small-molecule antiviral drugs like remdesivir or monoclonal antibody cocktails have shown only modest efficacy with moderately improved survival rates vs placebo in hospitalized patients^8,9^. Hence, there is still a great need for new and effective antiviral drugs against SARS-CoV-2.

SARS-CoV-2, like other betacoronaviruses, possesses multiple features that allow it to subvert our antiviral defenses and cause disease. SARS-CoV-2 codes for several structural and non-structural proteins that inhibit anti-viral signaling cascades^10,11^ as well as enzymes that cap and methylate (Cap1) its nascent transcripts^12,13^. By using a Cap1 structure, SARS-CoV-2 molecularly mimics our own messenger-RNA (mRNA), allowing it to escape recognition by the critical, antiviral RNA pattern recognition receptors (PRRs) IFIT1 and RIG-I^14–16^. Nevertheless, antiviral PRR activation is essential to the development of an effective immune defense, resulting in the release of type-I and type-III interferon (IFN), the antiviral state, direct antiviral effector functions in the infected cell, and the recruitment and formation of an appropriate adaptive immune response^16^. Thus, despite numerous escape mechanisms, type I IFNs still play a key role in SARS-CoV-2 immune defense^17,18^. A robust early type I IFN response is associated with the clearance of the infection whereas an absent or late induction of type I IFNs is associated with progression to severe COVID-19^18–20^. Moreover, anti-type I IFN autoantibodies or genetic alterations of the IFN pathway have been associated with severe COVID-19, underscoring the importance of this pathway for the anti-viral defense. ^21,22^

The use of synthetic nucleic acids to activate antiviral PRRs and induce the release of type-I IFN has been studied in numerous therapeutic contexts^23^. The cytosolic RNA receptor RIG-I has proven particularly suitable for such applications. RIG-I is activated by blunt-end double-stranded RNA with a 5’-tri- or 5’-diphosphate^24–26^, and potent specific ligands can be generated by solid phase synthesis^27^. Its activation induces a potent and broad antiviral program, which goes beyond the response induced by recombinant type I IFN alone^28^. Since RIG-I is widely expressed in nucleated cells including tumor cells, and its activation induces potent NK cell responses^29,30^, RIG-I ligands have been employed in the immune therapy against tumors in mouse models^31^, and in phase I/II clinical trials in humans^32^. Synthetic RIG-I ligands induce broad antiviral effects,^28^ and, in previous work, we showed that the stimulation of RIG-I was more effective than the stimulation of other nucleic-acid-sensing PRRs like TLR7 or TLR9 in protecting mice from a lethal influenza virus infection^33^.

In the current study, we investigated the efficacy of synthetic 5’triphosphate dsRNA RIG-I ligands (3pRNA) against SARS-CoV-2 in the murine K18-hACE2 mouse model of COVID-19. This mouse model recapitulates key aspects of human COVID-19 disease, such as anosmia, severe lung inflammation and impaired lung function^34–36^. Herein, we found that systemic RIG-I stimulation one to seven days prior to virus challenge substantially improves the clinical course of an otherwise lethal infection. Improved survival was associated with a lower viral load in the lungs and in the brain, demonstrating that RIG-I activation is a promising strategy for a short-term prophylactic treatment of SARS-CoV-2 and likely other emerging viral infections during the early phases of a pandemic, when vaccines are not yet available.

## Material and Methods

### Mice

B6.Cg-Tg(K18-ACE2)2Prlmn/J (K18-hACE2) were purchased from Jackson Laboratories and bred in-house at the University Hospital of Bonn. 8-to-20-week-old mice of both sexes were used throughout the study. All mice were housed in IVC cages in groups of up to five individuals per cage and maintained on a 12-hour light/dark cycle at 22–25°C temperature and 40–70% relative humidity under specific-pathogen free conditions. All mice were fed with regular rodent’s chow and sterilized water ad libitum. All procedures used in this study were performed with approval by the responsible animal welfare authority (81-02.04.2019.A433, LANUV NRW).

### Virus stock

The SARS-CoV-2 virus stock used in this study was raised from a throat swab isolate of a SARS-CoV-2-infected patient at the University of Bonn, Germany in March 2020 (SARS-CoV-2/human/Germany/Heinsberg-01/2020). The virus was passaged in VeroE6 cells and the viral titers were determined using a plaque assay as described in^37^.

### 3pRNA synthesis

Synthetic 3pRNA was chemically synthesized by solid-phase synthesis as described previously^27^. CA21-control RNA (5’-CACACACACACACACACACAC-3’) was purchased from Biomers.

### *In vivo* RIG-I stimulation

3pRNA or control RNA (CA21) was complexed with in vivo-jetPEI (N/P ratio 6) according to the manufacturer’s protocol (Polyplus). At the indicated time points, mice were injected intravenously with 20 μg RNA into the tail vein.

### SARS-CoV-2 mouse infection

All SARS-CoV-2 experiments were performed in a Biosafety Level 3 (BSL-3) facility at the University Hospital Bonn. K18-hACE2 transgenic mice were lightly anesthetized with ketamine/xylazine, before 5×10^4^ PFU of SARS-CoV-2 virus was pipetted onto the nose and subsequently inhaled by the animal. On day 1 to day 3, oral swabs were obtained using minitips (Copan, 360C) and placed in 1 ml UTM medium. Viral antigen in the oral swabs was quantified with ELISA according to the manufacturer’s protocol (SARS-CoV-2-Antigen-ELISA, Euroimmun). Following infection, weight loss and survival were monitored up to twice daily for 13 days. Endpoint criteria were >20% weight loss, lethargy, motor deficits and high respiratory rates.

### RNA isolation and qRT–PCR

Total RNA was extracted from mouse lung and brain tissues using TRIzol (Invitrogen) according to the manufacturer’s instructions. After a DNase I digestion step (ThermoFisherScientific), reverse transcription was carried out with RevertAid reverse transcriptase (ThermoFisherScientific) according to the manufacturer’s instructions. Viral RNA from oral swab material was purified using the NucleoSpin RNA Virus kit (Macherey & Nagel) according to the manufacturer’s instructions and used as a template for cDNA synthesis with RevertAid reverse transcriptase (ThermoFisherScientific).

The resulting cDNA was used for amplification of selected genes by qPCR using EvaGreen QPCR-mix II (ROX) (Biobudget) on a Quantstudio 5 machine (ThermoFisherScientific). SARS-CoV-2 Spike RNA expression was determined using the commercial E.Sarbecco primer sets (IDT, 10006888 and 10006890) with in 40 cycles measured. For tissues, relative expression values were normalized to murine *gapdh (fwd 5’* CTG CCC AGA ACA TCA TCC CT 3’, rev 5’ TCA TAC TTG GCA GGT TTC TCC A 3’), and average values from duplicates were presented as 2^-ΔCq^. Viral expression from oral swab material was presented as average values for each mouse as mean C_q_ values.

### IgG ELISA

The presence of SARS-CoV-2-specific IgG antibodies in the sera of mice was quantified using ELISA (QuantiVac, Euroimmun) according to the manufacturer’s recommendations, with the exception that the anti-human anti-IgG antibody in the kit was exchanged for a goat anti-mouse IgG-HRP conjugate (Jackson Immuno Research, 115-035-062), 1:5000 in 0.5% BSA in PBS.

### Histology

Half of the lungs and one mid-sagitally cut half of the brain were fixed in 6% neutral buffered formalin for at least 48 h. Tissues were embedded in paraffin. For SARS-CoV-2 antigen immunohistochemistry, slides were incubated with blocking reagent (10% normal goat serum) followed by rabbit monoclonal antibody against SARS-CoV-2 nucleocapsid protein (Biozol, SIN-40143-R019, 1:20000). The secondary antibody and the chromogen from the Dako REAL detection system (Agilent Technologies) was used for the staining according to the manufacturer’s protocol. Tissue sections were visualized using an Aperio SlideScanner CS2 and the Aperio Imagescope 12.4 software (Leica). Three scientists scored the sections in a blinded fashion as follows: 0 no staining; 1 weak staining, <5% brain area or <10% of lung area; 2 strong staining, 5-33% brain area or 10-50% of lung area; 3 strong staining >33% of brain area or >50% lung area.

### Statistical analysis

All statistical tests were calculated using Prism 9 (GraphPad). A p-value of <0.05 was considered statistically significant. Survival curves were analyzed using the log rank Mantel-Cox test. Analysis of weight change was determined by two-way analysis of variance (ANOVA). All results are expressed as mean + SEM and were corrected for multiple comparisons. Data were analyzed using a non-parametric Kruskal-Wallis test.

## Results

### Prophylactic treatment with RIG-I ligand protects K18-hACE-2 mice from a lethal challenge with SARS-CoV-2

K18-hACE2 mice overexpress the human ACE2 gene under the control of the Keratin18 promotor. In contrast to wildtype mice, SARS-CoV-2 replicates in K18-hACE2 mice leading to a lethal disease that has key characteristics of human COVID-19^34^. Compared to other animal models of SARS-CoV-2 infection, K18-hACE2 mice show the most severe phenotype and the highest lethality^38^.

To investigate the protective effects of a selective RIG-I ligand (3pRNA), K18-hACE2 mice were injected intravenously with a single dose of 20 μg 3pRNA complexed with *in vivo* jetPEI at seven days(d-7), three days(d-3) or one day(d-1) prior to inoculation with 5×10^4^ PFU SARS-CoV-2 (SARS-CoV-2/human/Germany/Heinsberg-01/2020). Weight loss and clinical manifestations were recorded daily for up to 13 days post-infection(dpi). Mice were euthanized for ethical reasons when they lost more than 20% of their initial body weight or showed overt signs of illness (Fig. 1A).

**Figure 1:**
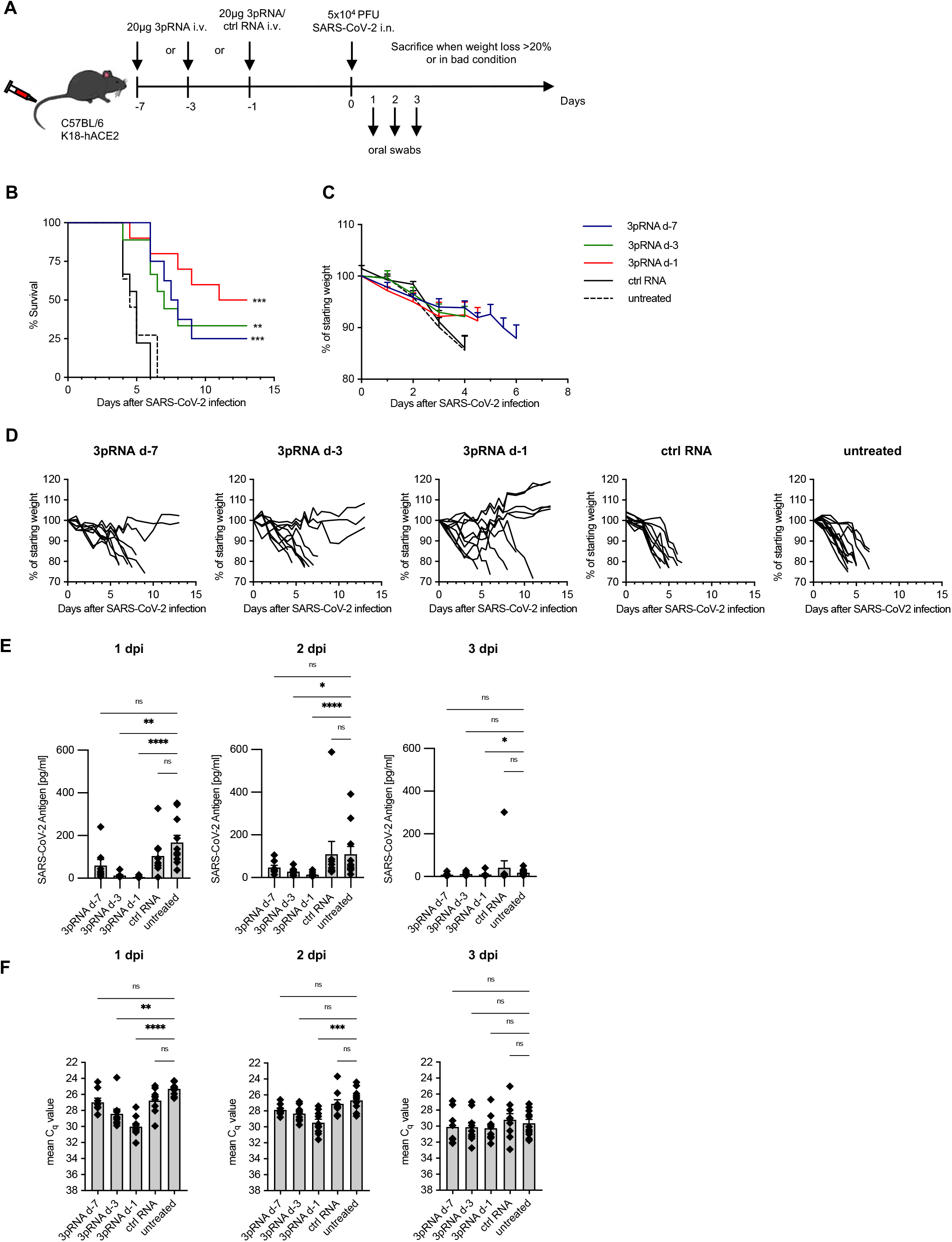
Prophylactic RIG-I stimulation protects mice from lethal SARS-CoV-2 infections. (A) Experimental setup. K18-hACE2^wt/tg^ mice were i.v. injected with 20 μg 3pRNA or control RNA complexed to *in vivo*-JetPEI on the indicated days. On day 0, mice were infected intranasally with 5×10^4^ PFU SARS-CoV-2. Oral swabs were obtained on day 1 to day 3 post-infection. Disease development and survival were monitored up to twice daily until 13 dpi. (B) Kaplan–Maier curve and (C) weight loss (pooled) of SARS-CoV-2-infected animals (D) Individual weight loss development of each mouse until their individual time of death or 13 dpi. (E+F) SARS-CoV-2 antigen ELISA (E) and expression of SARS-CoV-2 viral RNA (F) in oral-swab material on 1–3 days post-infection (dpi). Plotted are the mean + SEM (3pRNA d-7 n=8, 3pRNA d-3 & ctrl RNA n=9, 3pRNA d-1 n=10, untreated n=11). Data are pooled from two independent experiments. Statistical significance was calculated by log-rank Mantel–Cox test (B) and non-parametric one-way ANOVA (Kruskal–Wallis test) and Dunn’s multiple testing (E,F). * p<0.05, ** p<0.01, *** p<0.001, **** p<0.0001.

SARS-CoV-2-infected mice rapidly and uniformly showed weight loss and signs of disease such as reduced activity, hunched posture and lethargy. They all reached the euthanasia criteria by day 4.5 to 6 post-infection when left untreated (Fig. 1B-D). Some mice exhibited severe intestinal symptoms like bowel obstructions or neurological symptoms marked by progressive motor deficits. A single injection of 3pRNA administered one day (d-1) prior to inoculation with the virus improved the survival rate from 0% (treatment with control RNA) to 50% (Fig. 1B). Injection at three days (d-3) or seven days (d-7) days prior to infection still improved the survival rate by 25-30%. Some of the 3pRNA-treated mice that eventually succumbed to SARS-CoV-2 infection still showed slower weight loss and a delayed onset of other symptoms resulting in a right shift of the Kaplan-Meier curve (Fig. 1D). Surviving animals maintained or increased their weight and did not show any signs of disease for the duration of the experiment.

To monitor viral replication, oral swabs were taken one-to-three days post-infection. SARS-CoV-2 viral antigens were quantified by ELISA (Fig. 1E) and viral RNA was quantified by qPCR (Fig. 1F). Pretreatment with 3pRNA one (d-1) and three (d-3) days (with a trend for d-7) prior to infection led to a significant reduction of viral burden on day 1 and day 2 after viral challenge (Fig. 1E and F). In contrast, pre-treatment with control RNA had no effect compared to untreated animals. At 3 days after viral challenge, viral antigen was no longer detectable in most of the mice (Fig. 1E), suggesting the infection had been cleared from the upper respiratory tract where the swabs were taken.

### Mice prophylactically treated with RIG-I ligand demonstrate less viral spread in the lungs and brain

In line with the lower viral burden in the upper respiratory tract, we observed a significant decrease in viral RNA expression levels in the lungs of mice treated with 3pRNA one day before viral challenge (Fig. 2A). Notably, organs were collected on the day of death of each individual mouse. Thus, the endpoint of 3pRNA-treated animals was taken much later than untreated and control animals because of the prolonged survival of RIG-I ligand-treated mice. Surviving animals (marked in green) showed the most profound effect with complete clearance of viral RNA from the lungs.

**Figure 2:**
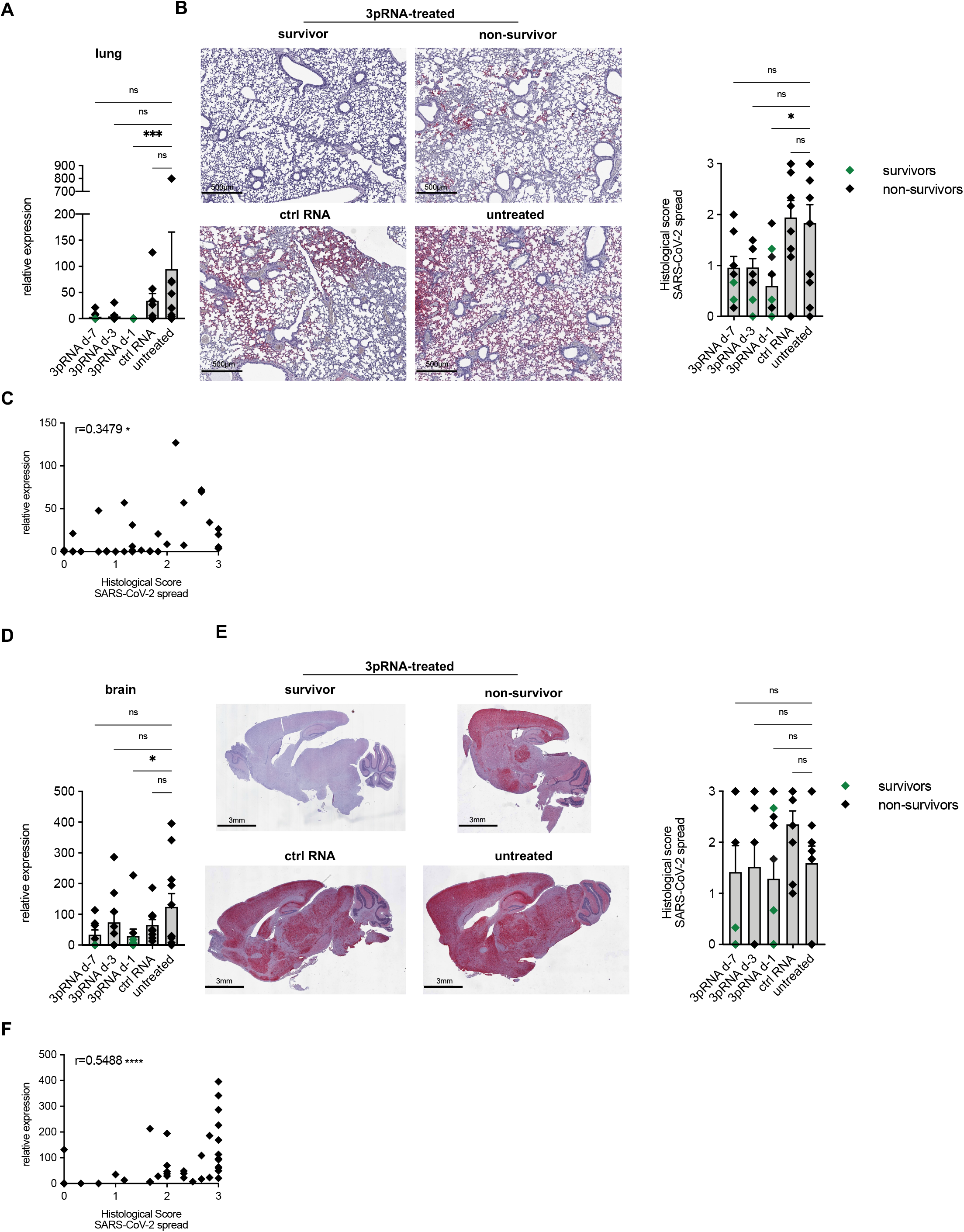
RIG-I prophylaxis reduces viral manifestation into the lungs and established adaptive immunity in surviving animals. (A) SARS-CoV-2 viral burden in the lungs quantified by qPCR relative to murine *gapdh* expression. (B) Representative pictures and scoring of immunohistochemical staining of the SARS-CoV-2 nucleocapsid protein in lung sections. (C) Correlation of histological scores of SARS-CoV-2 spread (x-axis) and relative expression qPCR values of SARS-CoV-2 spike expression in lungs. A normal distribution of values was assumed and for statistical analysis the Pearson coefficient was calculated as indicated (n=47). (D) SARS-CoV-2 viral burden in the brain quantified by qPCR relative to murine *gapdh* expression. (E) Representative pictures and scoring of immunohistochemical staining of the SARS-CoV-2 nucleocapsid protein in brain sections. (F) Correlation of histological scores of SARS-CoV-2 spread (x-axis) and relative expression qPCR values of SARS-CoV-2 spike expression for brain. A normal distribution of values was assumed and for statistical analysis the Pearson coefficient was calculated as indicated (n=47). Plotted are the mean + SEM (in A,B,D,E 3pRNA d-7 n=8, 3pRNA d-3 & ctrl RNA n=9, 3pRNA d-1 n=10, untreated n=11). Data are pooled from two independent experiments. Statistical significance was calculated by non-parametric one-way ANOVA (Kruskal–Wallis test) and Dunn’s multiple testing. * p<0.05, ** p<0.01, *** p<0.001, **** p<0.0001.

Immunohistochemical staining of SARS-CoV-2 nucleocapsid in lung sections correlated with the results of qPCR analysis when scored by three independent scientists in a blinded fashion (Fig.2C, r=0.3479, p<0.05), with significantly lower scores for mice pre-treated 1 day prior to challenge (Fig. 2A, B). In general, untreated and control RNA-treated mice showed broad positive staining for the nucleocapsid throughout the tissue. In contrast, mice pre-treated with 3pRNA that required euthanasia during the experiment (non-survivors) only presented a few focal spots of positive staining throughout the lungs, and 3pRNA-treated mice surviving the infection (again represented in green) presented with no or low positive staining (Fig. 2B).

Spread of viral infection into the central nervous system has been observed in COVID-19 patients and also described in the K18-hACE2 mouse model^34,36,39^ Viral RNA expression levels (Fig. 2D) and immunohistochemical staining (Fig. 2E) correlated more closely in brain sections (Fig. 2F, r=0.5488, p<0.0001) than in lung tissue (Fig. 2E, r=0.3479, p<0.05). Immunohistochemical staining further revealed a broad distribution of the virus throughout all brain areas except the cerebellum (Fig. 2D). Both viral RNA expression levels and immunohistochemical staining demonstrated more interindividual heterogeneity in the brain as compared to the lung. This is in line with previous reports of incomplete penetrance to the brain^35^. 3pRNA prophylaxis one day before challenge with virus significantly reduced the viral burden quantified by qPCR. Viral RNA was at the detection limit in the brain of surviving mice. Even mice that received 3pRNA treatment but eventually succumbed to SARS-CoV-2 infection exhibited lower viral loads in the lungs (Fig. 3A, C). In contrast, viral loads in the brain did not differ (Fig. 3B, D), demonstrating that the decisive clinical manifestation separating survivors from non-survivors in this specific mouse model is the degree of infection of the brain.

**Figure 3:**
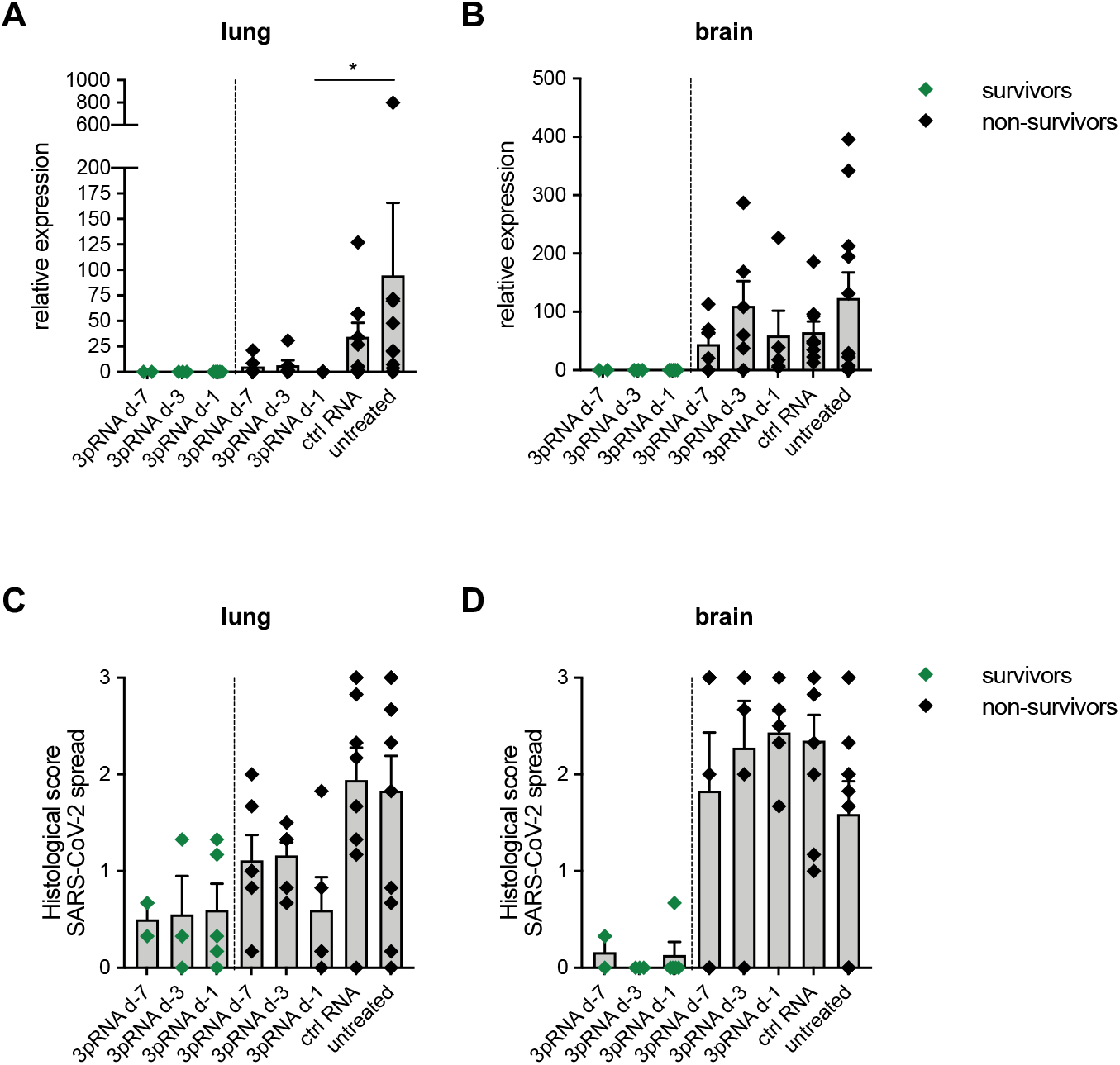
3pRNA-treated non-surviving mice demonstrated lower viral manifestation into the lungs. (A+B) SARS-CoV-2 viral burden in the lungs (A) and brain (B) quantified by qPCR relative to murine *gapdh* expression and separated by the survival of mice. (C+D) Scoring of the immunohistochemical staining of the SARS-CoV-2 nucleocapsid protein in lung (C) and brain (D) sections separated by the survival of mice. (surviving 3pRNA d-7 n=3, surviving 3pRNA d-3 n=3, surviving 3pRNA d-1 n=5, non-surviving 3pRNA d-7 & d-3 n=6, non-surviving 3pRNA d-1 n=5, ctrl RNA n=9, untreated n=11). Data are pooled from two independent experiments. Statistical significance was calculated by non-parametric one-way ANOVA (Kruskal–Wallis test) and Dunn’s multiple testing. * p<0.05, ** p<0.01, *** p<0.001, **** p<0.0001.

We then examined the levels of anti-SARS-CoV-2 specific IgG antibody titers in the sera of all surviving animals at the end of the observation period. While IgG titers of non-infected animals were below the detection limit, all 3pRNA-treated mice surviving the infection had substantial anti-SARS-CoV-2 specific IgG antibodies titers post infection (Fig. 4). These data demonstrate that the animals receiving 3pRNA prophylaxis successfully cleared the viral infection and established an adaptive immune response against SARS-CoV-2 as evidenced by the development of SARS-CoV-2 specific antibodies.

**Figure 4:**
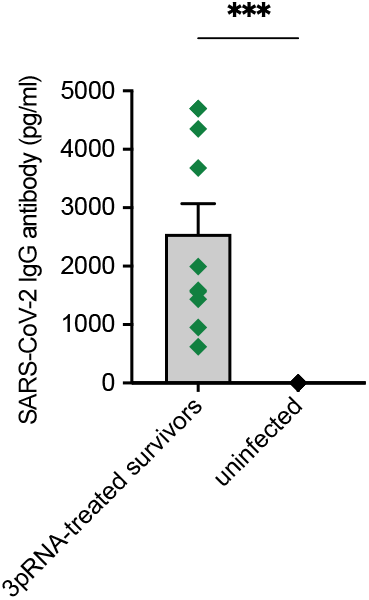
RIG-I prophylaxis established adaptive immunity in surviving animals. Quantification of anti-SARS-CoV-2 specific IgG antibody titers in sera of surviving animals (n=10 vs n=5 uninfected). Data are pooled from two independent experiments. Statistical significance was calculated by non-parametric one-way ANOVA (Kruskal–Wallis test) and Mann-Whitney-U test. *** p<0.001.

## Discussion

Innate antiviral defense is triggered upon immune sensing of viral nucleic acids. The betacoronavirus SARS-CoV-2 utilizes a number of mechanisms to efficiently escape from innate recognition of its viral RNA. In the present study, we evaluated whether prophylactic activation of innate antiviral immunity through i.v. administration of a specific RIG-I ligand protects mice from subsequent SARS-CoV-2 infection. We show that, in the murine K18-hACE2 mouse model of COVID19, synthetic RIG-I agonists drastically reduce viral replication not only in the upper and lower airways of SARS-CoV-2 infected mice but also in the central nervous system. Prophylactic treatment even up to seven days prior to infection substantially decreased morbidity and mortality compared to untreated animals. All RIG-I agonist-treated mice that survived the infection developed antibodies to SARS-CoV-2, showing that innate antiviral immunity acts in concert with adaptive immunity to provide protection against reinfection. It is important to note that the mouse model in this study exhibits 100% lethality, while for SARS-CoV-2 infection in humans the infection fatality rate is below 1%^40,41^. Thus, it is very likely that the results presented in this study largely underestimate the protective effect of prophylactic RIG-I ligand treatment against SARS-CoV-2 infection in humans.

Other studies have proposed stimulation of the innate immune system as an approach for SARS-CoV-2 treatment. Notably, recombinant PEG-IFNa2^42^, PEG-IFNb^43^ or PEG-interferon lambda^44^ treatment were shown to accelerate the recovery of COVID-19 patients in phase II trials. However, 3pRNA offers the additional advantage that the entire physiological spectrum of type I and III interferons are induced and that the anti-viral immune expression profile is broader than the one induced by type I IFN alone^28^. Moreover, treatment with recombinant interferon frequently leads to the generation of autoantibodies against the administered type I IFN which reduces the efficacy of the therapy^45^. In this context it is important to note that patients with autoantibodies against interferons have been reported to be more susceptible to severe COVID-19^22^. Simultaneous stimulation of different physiological forms of type I and III IFN by therapeutic RIG-I activation may have the advantage of inducing less anti-type I IFN autoantibodies. Moreover, the absence of anti-type I IFN autoantibodies is expected offer an advantage for future immune responses against other viral infections.

Consistent with our data, in a very recent pre-publication, Mao et al. have demonstrated efficacy of RIG-I agonists against SARS-CoV-2 infections^46^. While our study used 5’-triphosphate blunt-end double-stranded RNA, they utilized a 5’-triphosphate stem loop RNA. Nonetheless, both studies showed reduction in viral load and increased survival of K18-hACE2 mice with comparable doses of RIG-I agonist (20 vs 15 μg). However, it is important to note that, despite utilizing the same K18-hACE2 mouse model, these two studies differ in the virus dose used for infection and the resulting lethality. This needs to be considered when comparing quantitative efficacies of different compounds and prophylactic treatment regimens. Furthermore, in line with our previous findings on RIG-I-mediated protection against influenza virus infection^33^, Mao et al. confirmed that the efficacy of the RIG-I ligands depends on a functional type I IFN system.^33^

While vaccines will continue to be the most important weapon against SARS-CoV-2, the capability of RIG-I agonists to induce protection in immunocompromised hosts and their effectiveness against variants of concern^46^ would enable them to fill important niches for example by protecting transplantation patients. The reduced morbidity and mortality after prophylaxis even when administered seven days prior to infection in our study shows the induction of a long lasting anti-viral state by RIG-I agonists which would allow for a clinically feasible weekly pre-exposure prophylaxis in high exposure environments. Because double-stranded RIG-I agonists have already been tested in phase I/II clinical studies for oncologic indications (NCT03739138, NCT0306502)^32^, trials for the prophylaxis of COVID-19 could swiftly be initiated. Importantly, an effective antiviral prophylaxis of individuals exposed to newly emerging viruses, such as relatives of those infected or health care workers, may close the window for the development of a potential future pandemic early on before the virus becomes widespread and more difficult to contain.

## Acknowledgements

We thank Meghan Campbell for her critical reading of this manuscript and Hendrik Streeck for the use of the BSL3 facility at the Institute of Virology. We thank Patrick Müller, Sandra Ferring-Schmitt, and Janett Wieseler for expert technical assistance.

This study was funded by the Deutsche Forschungsgemeinschaft (DFG, German Research Foundation) under Germany’s Excellence Strategy–EXC2151–390873048 of which E.B., G.H., H.K. and M.S. are members. It was also supported by the Deutsche Forschungsgemeinschaft (DFG, German Research Foundation) – Project-ID 369799452 – TRR237 to E.B., G.H., B.M.K., H.K. and M.S. C.G. is the recipient of a BONFOR scholarship (University of Bonn). M.R. is funded by the Deutsche Krebshilfe through a Mildred Scheel Nachwuchszentrum Grant (Grant number 70113307).

